## Supplementary figures and images for "Phosphoproteomic identification of Mos-MAPK targets in meiotic cell cycle and asymmetric oocyte divisions"

### Source Data File 4

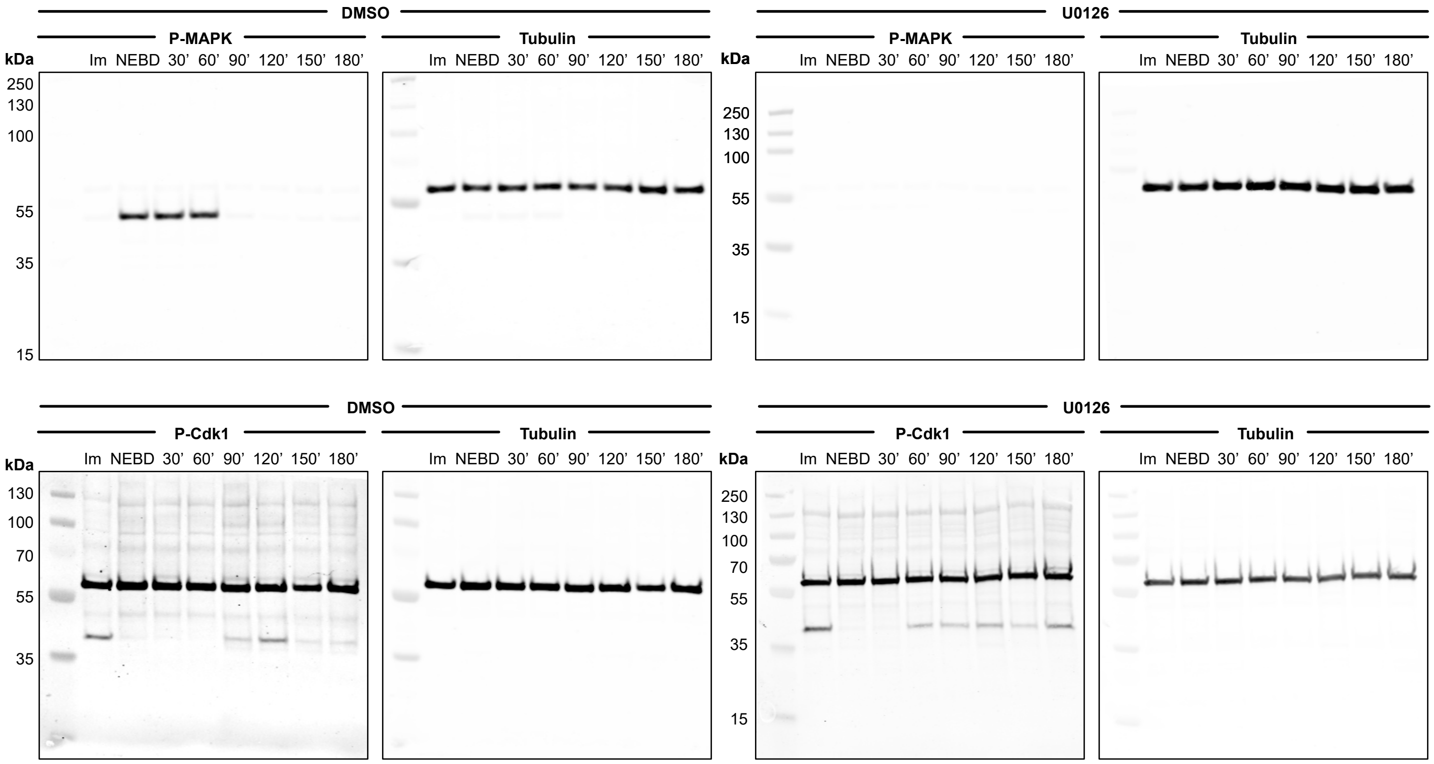

### Source Data File 5

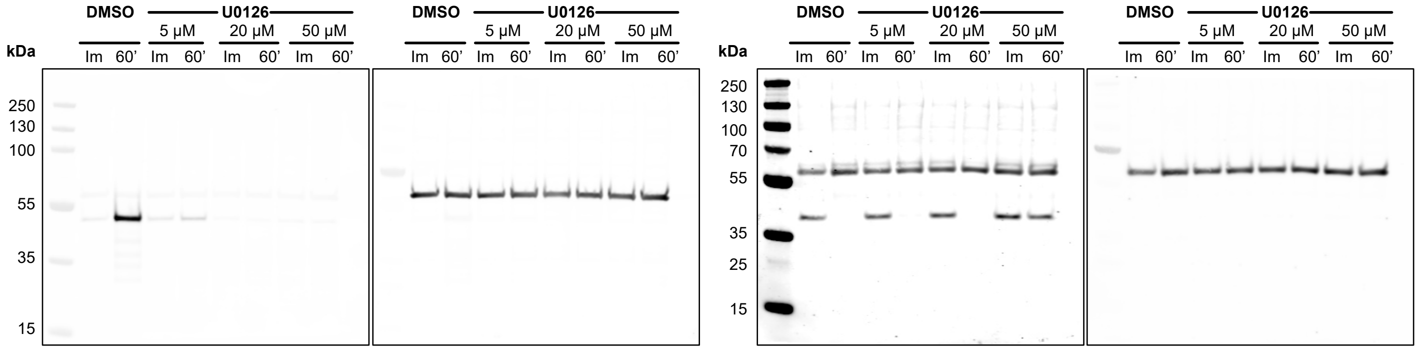

### Supplemental Figure 1

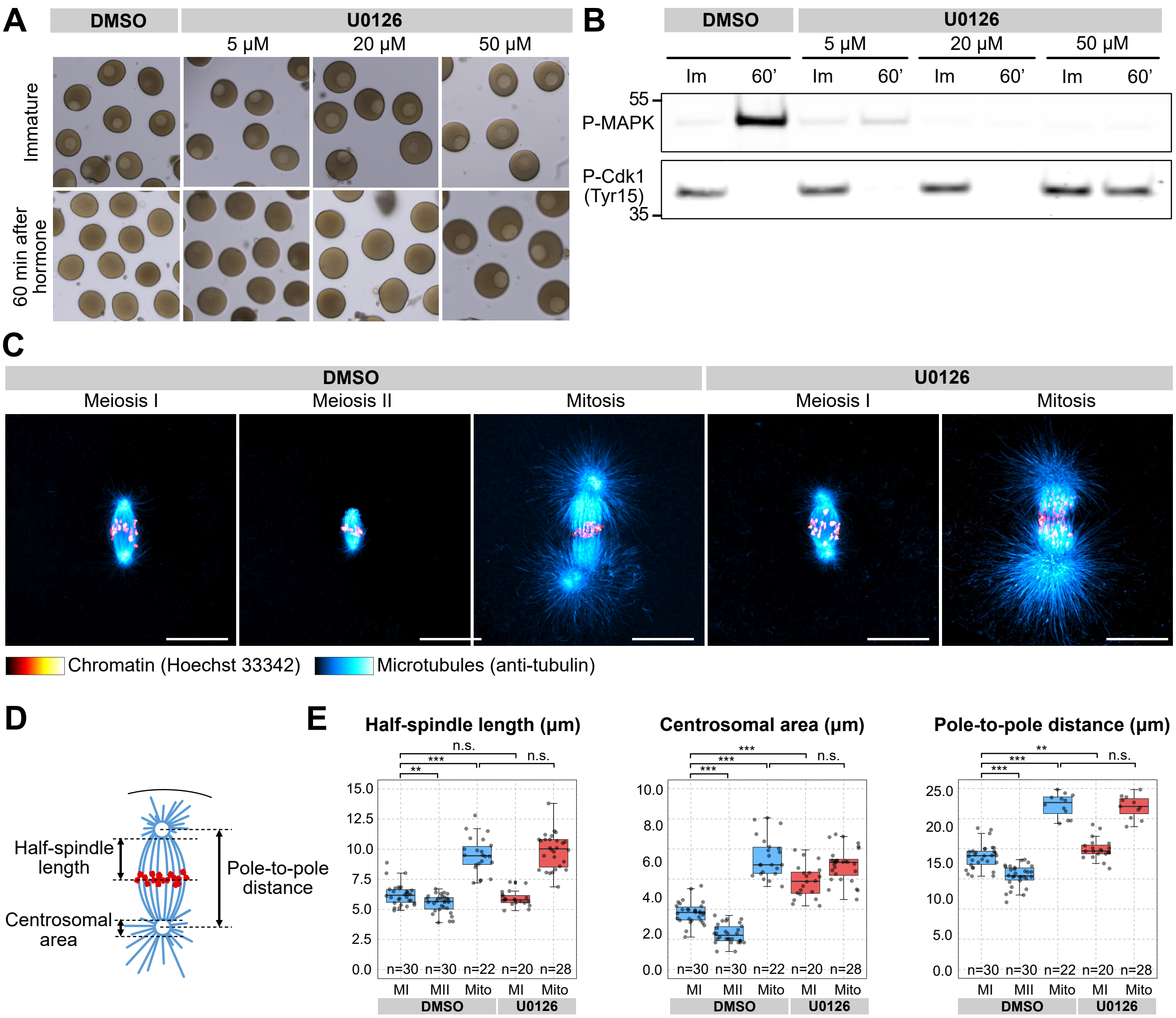

### Supplemental Figure 2

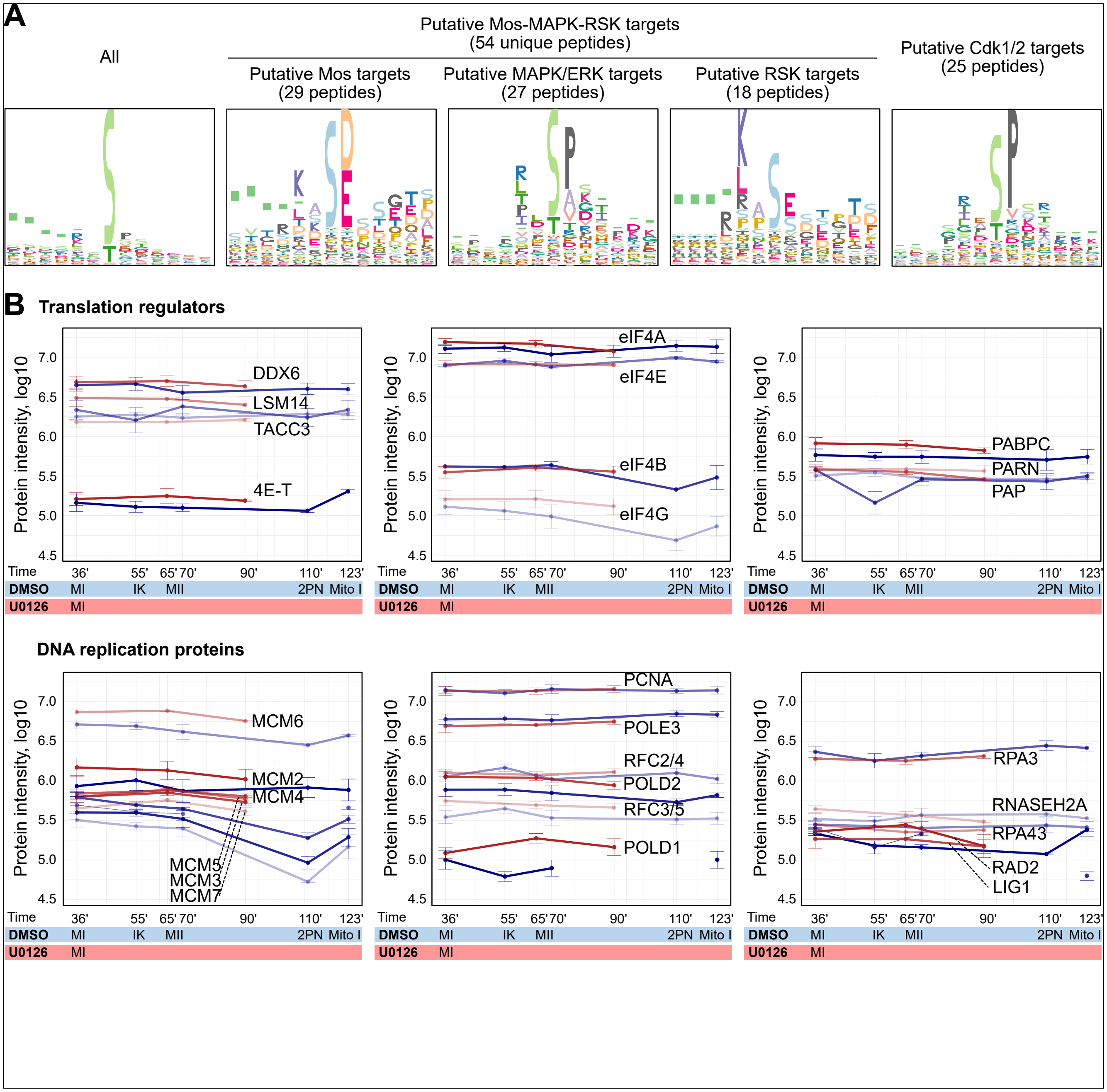

### Supplemental Figure 3

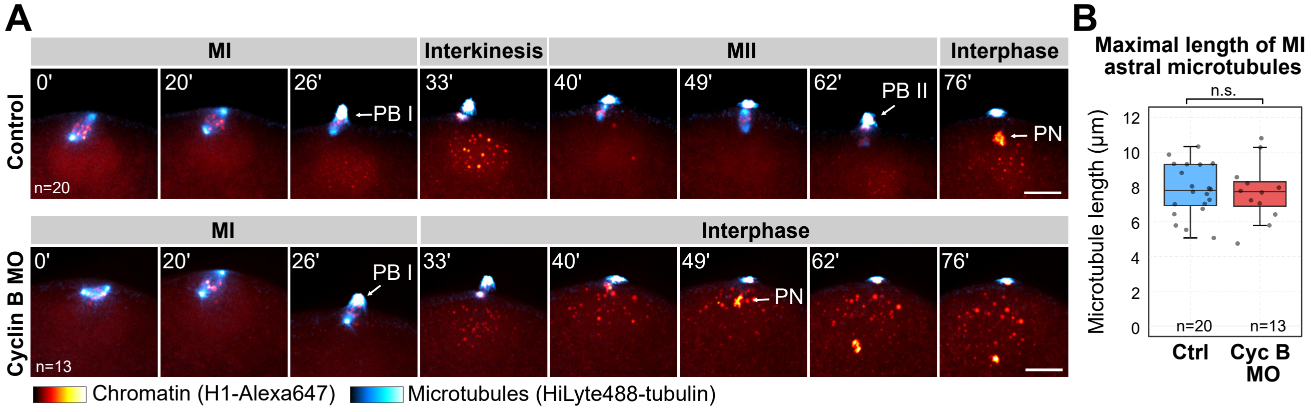
