## Supplemental Table 1 for "Phosphoproteomic identification of Mos-MAPK targets in meiotic cell cycle and asymmetric oocyte divisions"

### Table S1.

| | Name | KEGG | Protein IDs | Peptide | MS scan IDs | p-adj | Enrichment |
|---|---|---|---|---|---|---|---|
| **MAPK module** | MEK1 | K04369 | XP_038079443.1, PMI_017007 | NLTLPVKPDDSAPSNAVNNSS*ASMAAIAK | 51401 | 0.0444 | -1.5136 |
| | p90RSK | K04373 | XP_038049299.1, PMI_022459 | T*PKDSPGLPPSASAHQLFR | 44589 | 0.0165 | -1.6801 |
| **cdk1-Cyclin B module** | PPP2R2 | K04354 | XP_038070796.1, PMI_020469 | KRPLSTSDGT*AK | 22877 | 0.0180 | -1.5714 |
| | PPP2R2 | K04354 | XP_038070796.1, PMI_020469 | RPLS*TSDGTAKK | 23882 | 0.0040 | -1.8997 |
| | MYT1 | K06633 | PMI_008112 | AVSFRQSEPSVLQS*PHYNEK | 41151 | 0.0109 | -1.8548 |
| | APC3 | K03350 | PMI_002534, XP_038047876.1, PMI_014580 | LFS*NNSVKENATK | 45583 | 0.0003 | -2.4236 |
| | APC3 | K03350 | PMI_002534, XP_038047876.1, PMI_014580 | LFS*NNS*VK | 49005, 45078 | 0.0003 | -2.4422 |
| | CDC25C | K05866 | XP_038070722.1 | NRS*ETIMWEACNDKENVDIQNK | 42276 | 0.0262 | -1.4918 |
| | ARPP19 | | XP_038072705.1, PMI_013584 | KQS*TEISK | 23112 | 0.0004 | -2.3112 |
| **Translational activation module** | LARP1 | K18757 | PMI_014589 | EGRES*VDSPR | 8801, 7715 | 0.0000 | -3.9525 |
| | CPEB | K02602 | XP_038045408.1, PMI_005839, PMI_001490 | HTS*NNPGRPEK | 10213 | 0.0000 | -3.3840 |
| | CPEB | K02602 | XP_038045408.1, PMI_005839, PMI_001490 | YPS*QEIQQDYEK | 37277 | 0.0035 | -1.8924 |
| | EIF4ENIF1 | K18728 | PMI_009884 | KS*EPDGESGEKEEEGGDK | 21052, 19560 | 0.0000 | -2.9642 |
| | EIF3A | K03254 | PMI_009181 | S*AGIKDEEDEPR | 22666 | 0.0140 | -1.6728 |
| | EIF4B | K03258 | XP_038044282.1 | SNEDDSAFHKEEPLSPTSPTAPKS*PK | 36461 | 0.0440 | 1.5003 |
| | EIF4G1 | K03260 | XP_038075416.1 | VIQRVS*QTIK | 36609 | 0.0444 | -1.4466 |
| | GLD2/PAP | K14376 | XP_038074837.1, PMI_002048 | LPS*GELPDMSSPMPK | 53218, 49713, 47545 | 0.0111 | -1.7180 |
| | PPM1G | K17499 | PMI_005456, PMI_016115 | KAS*ESTPTDDDDSKR | 16212, 19904, 17237 | 0.0011 | -2.2632 |
| **Microtubule module** | CLIP1 | K10421 | XP_038055294.1, PMI_010288 | KTS*TSTVNSETSQR | 14508 | 0.0001 | -2.6735 |
| | CLIP1 | K10421 | XP_038055294.1, PMI_010288 | S*VDLTGNKPSLTNK | 42275 | 0.0232 | -1.5252 |
| | XMAP215 | K16803 | XP_038047612.1, PMI_009212 | SNRLS*QGSMSSEPVNGSAEQER | 22932 | 0.0107 | -1.7741 |
| | GTSE1 | K10129 | XP_038054192.1 | KES*DESQGSDSQESK | 15744 | 0.0003 | -2.4737 |
| | GTSE1 | K10129 | XP_038054192.1 | LPSTPST*PVHQDKK | 33139, 34006 | 0.0038 | -1.8905 |
| | GTSE1 | K10129 | XP_038054192.1 | LLSSDQS*ATKPVGR | 38688 | 0.0113 | -2.0316 |
| | POC1 | K16482 | XP_038054221.1, PMI_022968, PMI_011690 | S*TGADINAHSEPK | 21180 | 0.0022 | -2.0017 |
| | PCM1 | K16537 | XP_038056285.1, XP_038056290.1 | LLS*VQQQLR | 46891 | 0.0011 | -2.1149 |
| | CEP44 | K16761 | XP_038055484.1 | HAS*GSLVR | 15956 | 0.0211 | -1.5896 |
| | CEP44 | K16761 | XP_038055484.1 | RVS*VSVNELR | 34019 | 0.0482 | -1.3946 |
| | CEP192 | K16725 | XP_038068323.1 | RPS*FGTGHIKT*PEGR | 27259 | 0.0172 | -1.5904 |
| | KIF21 | K10395 | XP_038053367.1, XP_038053368.1 | LHSHSDRENES*ADEKEDAAEK | 18719 | 0.0460 | 1.5649 |
| | KIF14 | K17915 | PMI_006555, XP_038045418.1 | IDS*PMT*PLKR | 40449 | 0.0344 | -1.4586 |
| | FRYL | | XP_038055332.1, PMI_004592, PMI_010187 | RS*S*SGGTLEK | 17295 | 0.0113 | -1.6849 |
| **Cortical contraction module** | MYPT1 | K06270 | XP_038073240.1, XP_038073245.1 | TGS*ASTDTSSTSVR | 12906 | 0.0023 | -1.9539 |
| | MRCK | K16307 | XP_038071518.1 | RGGGS*VGAENNEVK | 17206 | 0.0032 | -1.9099 |
| | MRCK | K16307 | XP_038071518.1 | RLES*EKNTLSHR | 21479 | 0.0278 | -1.5109 |
| | mDia2 | K16688 | XP_038063096.1, PMI_026018, PMI_028305 | LQSRPS*VEEEEAHQEYER | 26257 | 0.0146 | -1.6930 |
| | MYO9 | K10360 | XP_038073351.1, PMI_008384 | KQS*DPQQAAEALGLPSDKPGK | 42738 | 0.0337 | -1.5479 |
| | BCR | K08878 | XP_038059104.1 | LS*TPDVEILNVR | 53153 | 0.0480 | -1.4272 |
